# *Dici*: a novel DNA transposon reshaping the genome of the opportunistic yeast *Diutina catenulata*

**DOI:** 10.1101/2025.11.06.686975

**Authors:** Frédéric Bigey, Marc Wessner, Martine Pradal, Corinne Cruaud, Jean-Marc Aury, Cécile Neuveglise

## Abstract

*Diutina catenulata* is an ascomycetous yeast of both environmental and emerging clinical relevance. However, genetic and genomic information on this species remains limited. In this study, we present a series of genomic analyses of selected *D. catenulata* strains, encompassing the full spectrum of their phylogenetic diversity. The species exhibits pronounced genetic structuring, with distinct clades representing deeply diverged lineages that show limited gene flow. This phenomenon is likely the result of clonal propagation or historical divergence. This study also uncovers a wide range of genes in both dairy and clinical isolates that are homologous to the drug resistance and virulence factors found in *Candida albicans*. A notable feature of *D. catenulata* genomes is an extensive degree of chromosomal instability associated with a novel DNA transposon family, called *Dici*, which has invaded all lineages. The high sequence conservation of *Dici* copies suggests a recent and potentially ongoing transpositional burst that is actively reshaping genome architecture. While the primary drivers of clade diversification remain to be elucidated, transposon activity and chromosomal rearrangements may have contributed to reproductive isolation and maintained genome plasticity. This multilayered genomic landscape provides a valuable model for studying DNA transposon proliferation, genome evolution, adaptation, and the emergence of pathogenicity in yeasts.

## 1 Introduction

Transposable elements (TEs) are DNA sequences capable of moving within and between genomes, significantly shaping genome architecture and evolution. They are broadly classified into two major classes based on their transposition mechanisms [1, 2]. Class I elements (retrotransposons), which transpose via an RNA intermediate through a “copy-and-paste” mechanism, are subdivided into long terminal repeat (LTR) elements, long interspersed nuclear elements (LINE), and short interspersed nuclear elements (SINE). For example, *Ty3* /*gypsy* (Metaviridae) and *Ty1* /*copia* (Pseudoviridae) are two major superfamilies of LTR retrotransposons that occur in all major groups of eukaryotes. In some large-genome plants, RNA transposons make up the majority of the nuclear genome like in maize (*Zea mays*) and wheat (*Triticum aestivum*) [3]. Class II elements (DNA transposons), which typically move via a “cut-and-paste” mechanism with a DNA intermediate, include four major groups: DDE transposases, Cryptons, Helitrons and Mavericks/Polintons. DDE transposons are the most diverse and widespread, with large superfamilies defined by phylogenetically distinct transposases [2]. Notably, the *IS630-Tc1-mariner* group of DDE transposons is broadly distributed across eukaryotes [4].

Yeast are unicellular fungi that reproduce asexually primarily through budding or fission. They harbor a variety of TEs that have contributed to genome diversification, gene regulation, and adaptation to different environments. In *Saccharomyces cerevisiae*, TEs are predominantly represented by LTR-retrotransposons named *Ty* elements, which resemble retroviruses in structure and replication mechanism [5]. The abundance and activity of these elements vary among yeast species and strains, reflecting both phylogenetic divergence and ecological pressures [6]. Some yeasts also possess DNA transposons, although these are less common and often degenerated [7, 8]. Through insertional mutagenesis, homologous recombination between TE copies, and the generation of chromosomal rearrangements, TEs actively promote genome plasticity in yeasts [9, 10]. Although TEs drive genetic innovation and adaptation, their uncontrolled activity can threaten genome integrity by inducing insertional mutations, chromosomal rearrangements, and genome size expansion [11, 12]. To mitigate these risks, host organisms have evolved sophisticated regulatory mechanisms, including RNA interference, epigenetic silencing [13], and domestication of TE-derived sequences [14]. An interesting example in yeast is the domestication of two distinct DNA transposons for mating-type switching in *Kluyveromyces lactis*: *α3* with homology to the transposase of mutator-like elements (MULE) and *KAT1* to *hAT* transposases [15, 16].

*Diutina catenulata* (formerly *Candida catenulata*) is an ascomycetous yeast belonging to the taxonomic order of Serinales (also known as the CUG-Ser1 clade) and to the Debaryomycetaceae family [17, 18]. This species is predominantly associated with environmental and food-related niches, particularly dairy products such as cheeses [19–23], where it contributes to the yeast diversity of various French cheeses [24–27]. Its prevalence in the gastrointestinal tract of poultry [28] and fish [29] has also been well documented, as well as its occurrence in the feces of pigeons [30].

In addition, *D. catenulata* has been identified in clinical specimens, including blood and mucosal surfaces. It has the capacity to colonize the digestive tract and trigger both superficial and deep infections [31–33]. These findings underscore its emerging pathogenic potential in immunocompromised individuals. Despite limited reports of human infection by *D. catenulata*, the emergence of multidrug resistance, particularly against frontline antifungals (azoles and echinocandins), raises significant concerns for clinical management should invasive infections arise [33, 34]. The dual ecological role of *D. catenulata* underscores the importance of understanding its genetic diversity, adaptive strategies, and potential impact on food safety and human health.

To date, only two studies have characterized the *D. catenulata* genome. O’Brien et al. [35] produced the first draft genome assembly from a soil isolate (accession: PJEZ01000000), thus confirming its placement within the Debaryomycetaceae family. Subsequently, Boden et al. [36] reported the genome assembly of the type strain CBS 565^T^ (accession: JBKQXV010000000), identifying virulence factors commonly found in *Candida albicans*. However, although genome annotation was reportedly performed in both studies, the available datasets contain only raw genome sequences, limiting their utility.

In the present study, we assessed the genetic diversity and population structure of *D. catenulata*. High-quality genome assemblies and annotations were generated for five representative strains spanning the species’ phylogeny. We provide evidence of a genome invasion by a novel DNA transposon, named *Dici*, which defines a previously unrecognized family in the *IS630-Tc1-mariner* group and likely contributes to the extensive chromosomal fragmentation observed in *D. catenulata*.

## 2 Results

### 2.1 Marked population structure in *D. catenulata*

A collection of 61 *D. catenulata* strains was assembled from French Protected Designation of Origin (PDO) cheeses [22] and from international yeast collections and laboratories, representing eight geographical and five ecological origins (Supplementary Table 1). Flow cytometry revealed that most strains are haploid (*n* = 51), including the type strain CBS 565^T^, while seven are diploid. Short Illumina sequencing reads were obtained for the 61 *D. catenulata* strains and were mapped to the assembled genome of strain CBS 565^T^ (see bellow). A total of 508,773 variants were identified, including 436,405 SNPs and 72,368 small insertions and deletions (indels). The overall nucleotide diversity (*π*, the average number of pairwise differences per nucleotide site) was estimated to be 6.2 *×* 10^−3^. This value is close to the values obtained in other yeast species associated with anthropized environments, such as *S. cerevisiae* (*π* = 3 *×* 10^−3^) and *Lachancea thermotolerans* (*π* = 9.3 *×* 10^−3^), while lower than that observed for *Kluyveromyces lactis* (*π* = 2.8 *×* 10^−2^) [37–39]. In the case of *D. catenulata*, the observed *π* value may be influenced by a sampling bias resulting from the fact that the majority of the strains analyzed were from French cheeses (*n* = 48) and by the limited number of clinical isolates (*n* = 9).

A high-quality subset of 351,942 SNPs was employed to construct a phylogenetic tree, which revealed six distinct clades, hereafter referred to as clade #1 through clade #6 (Fig. 1). The population structure was analyzed using a pruned subset of 34,101 SNPs that are in approximate linkage equilibrium with each other (*r*^2^ *<* 0.5) and the Bayesian model-based clustering approach implemented in fastStructure (Fig. 1). The analysis yielded a most probable population structure comprising six subpopulations, which correspond to the phylogenetic clades identified.

**Fig. 1.**
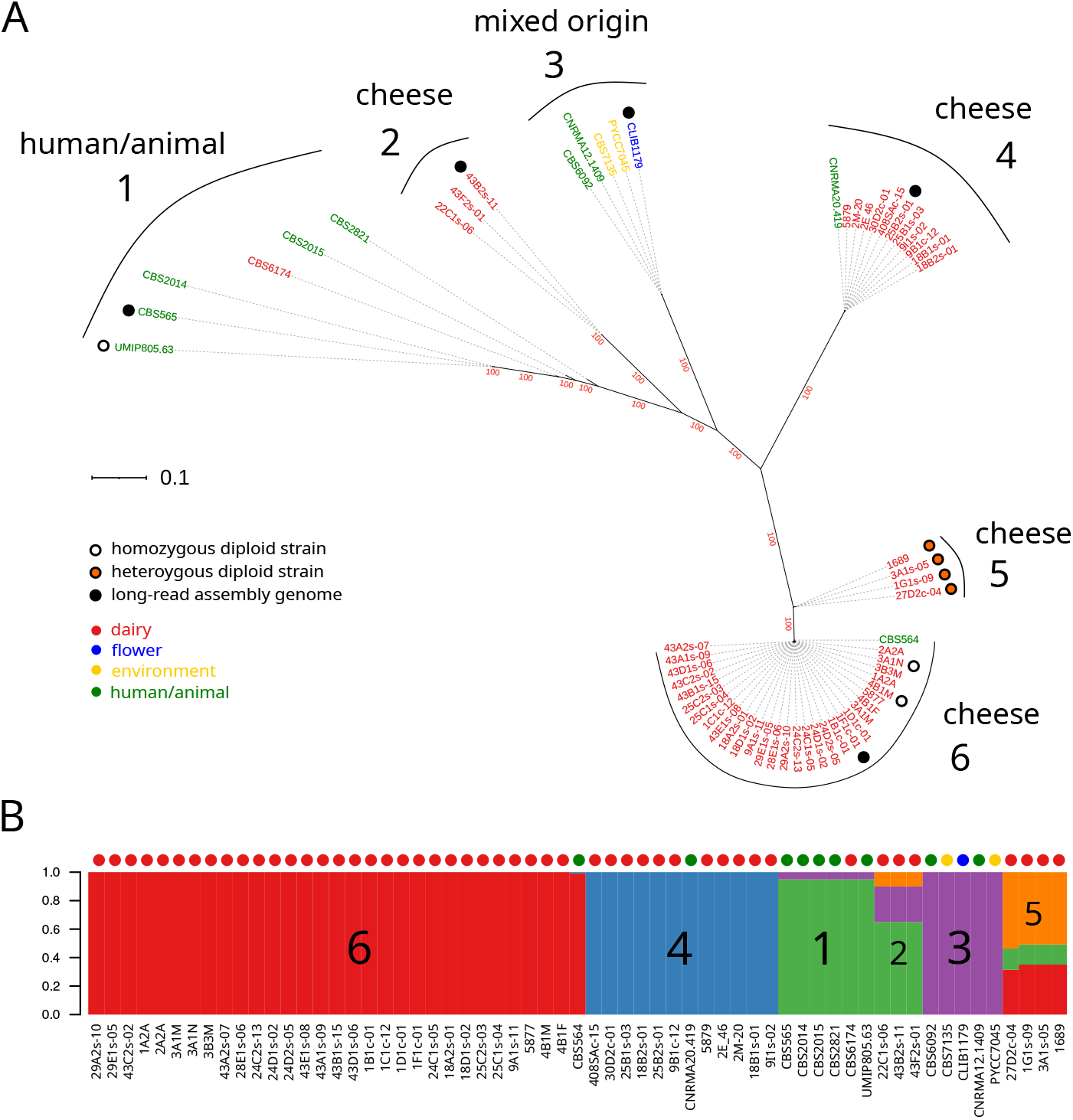
Phylogeny and population structure of the 61 *D. catenulata* strains. (A) maximum likelihood phylogenetic tree obtained with 351,942 SNPs (IQ-TREE, evolutionary model: TVM+F). Bootstrap values for the main branches are in red; (B) population structure obtained with a 34,101 SNPs subset in approximate linkage equilibrium (*r*^2^ *<* 0.5). The vertical axis shows the fractional representation of the resolved genetic groups (colors) within each strain (horizontal axis) for K assumed ancestral genetic groups. Model components used to explain the structure in the data *K* = 5

Clade #1, which exhibited the highest level of nucleotide diversity (*π* = 1.9*×*10^−3^), comprised six strains (UMIP805.63 was a homozygous diploid of strain CBS 565) isolated from human subjects, with the exception of CBS 6174, which was isolated from a milk sample obtained from a cow with mastitis. Clade #2 comprised three distinct cheese strains. Two of these strains were isolated from PDO #43, but were derived from two different producers (43B and 43F). These two strains differ by only 51 SNPs. The diversity within this group was the lowest observed (*π* = 7.1 *×* 10^−6^) across the clades. While clade #3 (*n* = 5) is the most diverse concerning its ecological origin (human, water, soil and flower), a low nucleotide diversity of 1.8 *×* 10^−5^ was observed, with less than 541 SNPs between strains. Clade #4 (*n* = 12; *π* = 3.4 *×* 10^−5^) is almost composed of cheese strains, with three couples isolated from the same PDO (#9, #18 and #25) and even from the same producers but at different seasons (18B1/18B2 and 25B1/25B2). Strain CNRMA20.419, isolated from a human hemoculture sample, was found to differ to no more than 851 SNPs with the cheese strains. The four heterozygous diploid strains of our collection, all isolated from cheese, presented high levels of heterozygosity (heterozygous SNPs/homozygous SNPs = 0.8) and were found to cluster together in clade #5. This clade displayed the highest diversity (*π* = 5.7 *×* 10^−3^), with strains differing by up to 9,103 SNPs. The population structure analysis revealed that these strains exhibited a mosaic pattern, indicative of a genetic contribution from members of clade #6 (cheese), clade #1 (human/animal) and an as yet unidentified contributor (illustrated in orange on Fig. 1). The large clade #6 (*n* = 31; *π* = 7.2 *×* 10^−5^) was composed of strains isolated from cheese. Several were from the same PDO cheeses: e.g. PDO #43 (*n* = 6) and #24 (*n* = 4) but from different producers or at different seasons. This clade contains two homozygous diploid cheese strains. It also includes an isolate obtained from a case of stomatitis (CBS 564) that does not grow at 35 °C, suggesting that this strain is opportunistic. Interestingly, strains isolated from the same PDO were found in different clades: PDO #18 and #25 were found in clades #4 and #6, and PDO #43 in clades #2 and #6. In the latter case, strains 43B2s-11 and 43B1s-15 were isolated from cheeses produced by the same producer in two different seasons.

In order to validate the genetic structure of the yeast population, an Analysis of Molecular Variance (AMOVA) was performed based on the SNP data (Supplementary Table 2). The results indicated that a large proportion of the total genetic variance (93.6%) could be attributed to variations among the six predefined clades, while the variance among individuals within clades was found to be negligible. These findings indicate a robust genetic structure of the *D. catenulata* population, with clades representing well-differentiated lineages.

Pairwise fixation indices (*F*_*ST*_) supported this observation, with high levels of genetic differentiation observed between clades, consistent with long-term divergence and limited gene flow between groups (Supplementary Table 2). In addition, elevated Tajima’s *D* values (*D >* 0.2 for clades #4 and #6; *D >* 0.8 for the others) indicate a decrease in population size and/or balancing selection. When clades were analyzed jointly, Tajima’s *D* was still consistently positive (*D* = 1.83). This pattern is most likely explained by a strong population structure, which inflates estimates of intermediate-frequency polymorphisms.

These results highlight the high level of genetic differentiation among the clades of *D. catenulata*, with clades likely reproductively isolated, possibly due to clonal propagation and historical divergence. The cheese strains are distributed across four distinct clades (#2, #4, #5 and #6). The presence of two strains isolated in humans among the dairy clades #4 and #6 does not necessarily imply that these strains are pathogenic, but that they could occasionally be human commensal.

### 2.2 A DNA transposon has invaded the *D. catenulata* genome

Four *D. catenulata* haploid strains (43B2s-11, CLIB 1179, 25B2s-01 and 1F1c-01), in addition to the type strain CBS 565^T^, were selected from the above genetic clades (excluding clade #5, which contains only diploid strains). Their genomes were sequenced, assembled, and comprehensively annotated, providing a foundation for comparative genomic analyses. A review of the key genomic features is presented in the accompanying documents (Supplementary Information; Supplementary Table 3). Notably, this analysis revealed the presence of a broad repertoire of genes homologous to *C. albicans* drug resistance and virulence determinants (Supplementary Table 4). Several of these genes occurred in multiple copies. This finding suggests that *D. catenulata* possesses the ability to develop antifungal resistance and pathogenicity, despite its relatively lower clinical prevalence compared to *C. albicans*.

The main Class I transposable elements (retrotransposons) identified in the genomes belong to two superfamilies: *Ty3* /*gypsy* (Metaviridae), for which four families were found in *D. catenulata*, and *Ty5* /*copia* (Pseudoviridae), for which two families were detected (Supplementary Table 5). Together, these elements comprise a relatively low percentage of the genome of the 61 *D. catenulata* strains (Fig. 4), and are almost entirely absent from some strains (CNRMA20.419, CBS 2014 and CBS 2821). The *Ty5* /*copia* elements are the most abundant, being particularly prevalent in seven strains and accounting for up to 0.5% of their genome. LINE retrotransposons (*L1* superfamily) were detected, while they were found mainly as truncated or degenerate copies (Supplementary Table 5).

During the annotation curation of the five *D. catenulata* genomes, a repeated sequence containing three coding sequences (CDSs) was identified (Fig. 2). It is flanked by two terminal inverted repeats (TIRs) and a systematic duplication of TA dinucleotides was observed on each side, a consequence of a target site duplication (TSD) commonly found at the insertion sites of the transposons of the *IS630-Tc1-mariner* superfamily [40]. The CDS located at the center of the element was found to encode a transposase (see below). This putative transposable element (TE) was designated *Dici*, for *D. catenulata* invader. Two types of *Dici* were identified, designated *Dici1* and *Dici2*, with lengths of 4.0 kb and 4.2 kb, respectively. The two elements showed marked divergence, with 52.2% nucleotide identity, but shared an identical GC content of 42.7%, considerably lower than the 53.5% observed for the *D. catenulata* nuclear genome.

**Fig. 2.**
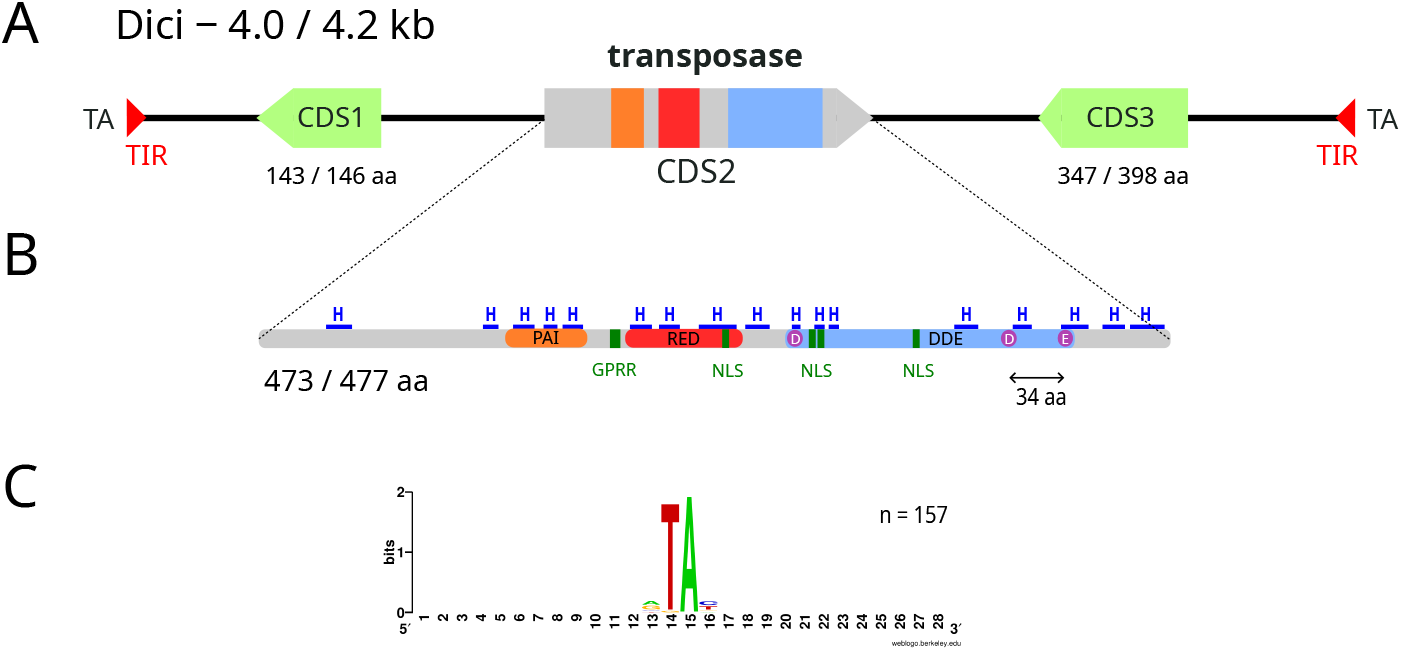
Characteristics of *Dici* DNA transposons detected in *D. catenulata*. (A) the 4-kb sequence is flanked by short terminal inverted repeats (TIRs), and a target site duplication (TSD) was observed at both extremities (TA). Three coding sequences (CDSs) are present. The transposase is located in the center of the element; (B) Schematic representation of the main characteristic features of the *Dici* transposases, namely a pair of helix-turn-helix motifs (PAIRED domain, orange and red regions), each consisting of three *α*-helices (H in blue). A conserved GRPR-like motif (also known as an AT-hook) is present between the two HTH motifs. Four putative nuclear localization signals (NLS), composed of basic amino acids (K or R), were detected in conserved regions. Finally, the catalytic triad DDE (highlighted with purple circles) is present within the catalytic domain (blue region), with 34 amino acids between the second aspartic acid (D) and glutamic acid (E). A complete alignment of the *Dici* transposases can be found in Supplementary Fig. 3; (C) Nucleotide composition at the insertion points was visualized using sequence logo. The number of sequences used is indicated

*Dici1* was predominantly identified in strain CBS 565 (96 full-length copies), while *Dici2* was found in the genomes of strains CLIB 1179, 43B2s-11, 25B2s-01 and 1F1c-01, with 8, 28, 39 and 45 full-length copies, respectively (Supplementary Table 6). Four cases of insertion of a *Dici1* transposon into another *Dici1* element were observed for strain CBS 565. Miniature inverted-repeat transposable elements (MITEs) derived from *Dici* were identified in the genomes of the five strains, with a length of 1,053 bp (Supplementary Table 6). TIRs of 8 and 33 bp, starting with the sequence CAGTCGGT, were detected at each extremity of *Dici1* and *Dici2*, respectively. Neither *Dici1* nor *Dici2* were detected in the genome of other *Diutina* species.

### 2.3 Massive genomic rearrangements in the *D. catenulata* genome

The five *D. catenulata* strains displayed pronounced variation in their karyotype profiles (Fig. 3), suggesting extensive chromosomal rearrangements. This was confirmed by whole-genome alignments, which revealed large-scale genome reshuffling between strains (Supplementary Fig. 1).

**Fig. 3.**
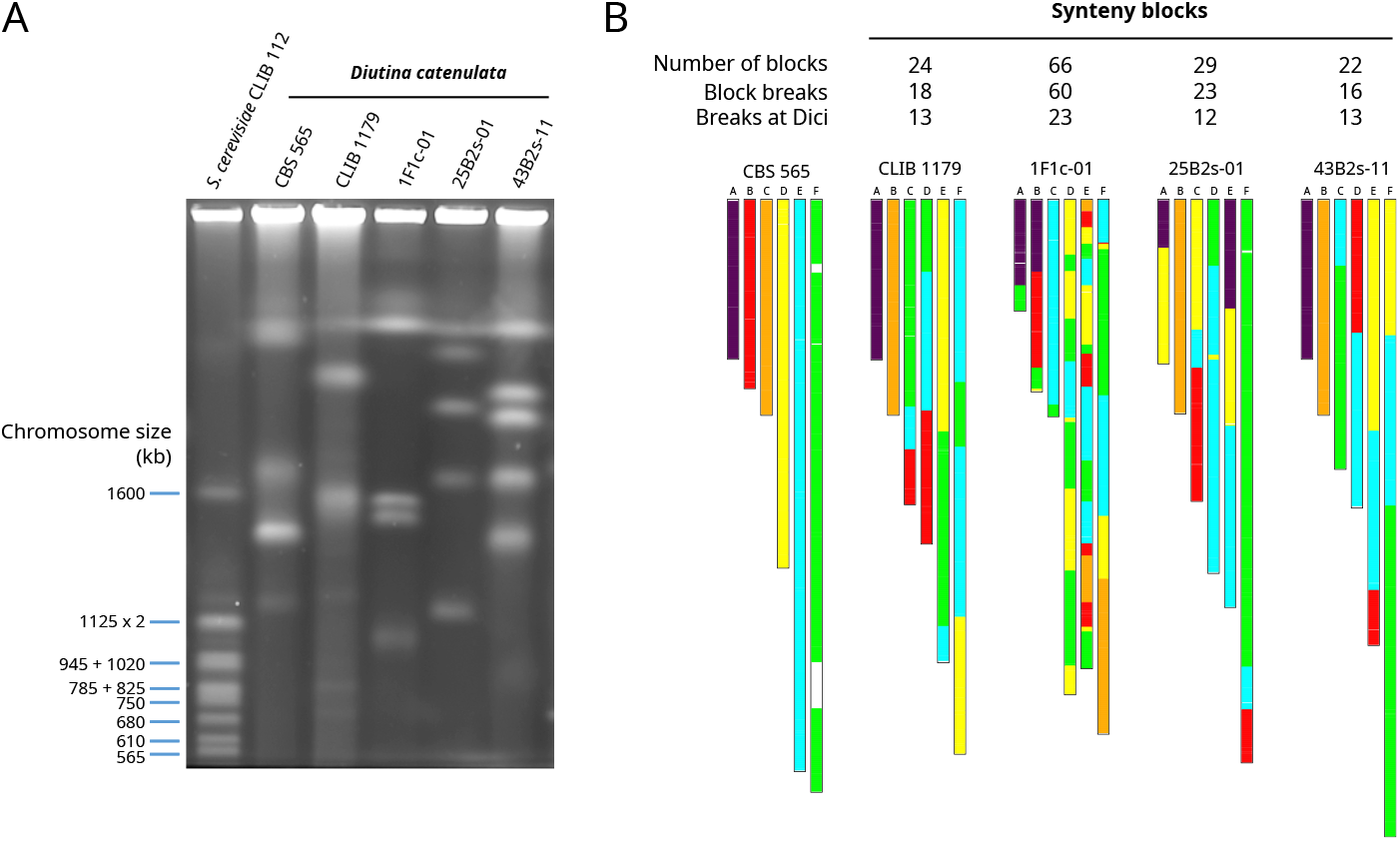
Chromosomal rearrangements in the *D. catenulata* genome. (A) karyotypes of the five *D. catenulata* strains exhibit distinct patterns of chromosome size. Numbers on the left represent the size of the *S. cerevisiae* chromosomes used as markers; (B) visualization of synteny blocks along the chromosomes. The six chromosomes of strain CBS 565 were used as references and given a color. The same color is used to show the corresponding synteny blocks in the other genomes. White blocks in the reference genome indicate regions of segmental duplication that were not used in the reconstruction of synteny blocks. The number of synteny blocks, the number of synteny block breakpoints, and the number of breakpoints at *Dici* element are indicated above the plot

Consistently, comparative analyses revealed a high number of synteny blocks, indicative of a lack of chromosome collinearity between strains (Fig. 3). To investigate the underlying causes of this genomic instability, we examined synteny breakpoints for associated sequence features. The borders of many synteny blocks were enriched in *Dici* transposons (Supplementary Table 7). The highest number of synteny blocks (*n* = 66) was observed in the comparison between strains CBS 565 (which harbors 96 copies of *Dici1*) and 1F1c-01 (45 copies of *Dici2*), where *Dici* elements were implicated in more than 20% of the observed breakpoints. These findings suggest a link between transposon copy number and the extent of chromosomal rearrangement, supporting the hypothesis that *Dici* activity contributes to genome structural plasticity in *D. catenulata*.

Thus, although genome plasticity enhances adaptive potential, the chromosomal rearrangements it generates may decrease hybrid fertility. This ultimately influences population structure, driving long-term divergence and reproductive isolation. This is consistent with the robust genetic structure observed in the *D. catenulata* population, suggesting that clades are likely to be reproductively isolated.

### 2.4 *Dici* is widely distributed in *D. catenulata*

We assessed the diversity of *Dici* elements across the *D. catenulata* assembled genomes (Supplementary Table 6). Within individuals, *Dici1* and *Dici2* were highly conserved. For instance, the 96 full-length copies of *Dici1* of CBS 565 exhibited an average pairwise nucleotide identity of 99.9%, while *Dici2* exhibited a comparable level of conservation, with identities ranging from 99.6 to 99.8% in strains 43B2s-11, CLIB 1179, 25B2s-01, and 1F1c-01. Likewise, the comparison of *Dici2* elements across strains (CBS 565 was excluded as it only contains two copies of *Dici2*) revealed extremely low levels of nucleotide divergence, with identities ranging from 99.4 to 99.6%. These values are more than 1% above those obtained from whole-genome comparisons, where identities ranging from 97.7 to 98.2% were observed when comparing the genomes of strains 43B2s-11, 1F1c-01, 25B2s-01 and CLIB 1179 (Supplementary Table 6). These results indicate that *Dici* elements were highly conserved within strains and exhibited limited diversification between clades. This finding suggests that *Dici* have undergone recent amplification and have exhibited minimal evolutionary divergence among clades.

An evaluation was conducted to determine the distribution of *Dici* elements across the 61 strains of our collection. The prevalence of *Dici* exhibited significant variation, with ranges extending from 0.03 to 2.70% of the genome (Fig. 4). The highest abundance was observed in clade #1 for strains CBS 565 and its sibling diploid UMIP805.63 (2.70%), both of which were found to be dominated by *Dici1*. In contrast, for the same clade, CBS 2821 exhibited the lowest *Dici* content (0.03%), while CBS 2014 was pre-dominantly dominated by *Dici2* (1.86%). Similarly, strains from clade #3 exhibited significant variability in *Dici* abundance. For instance, clinical isolates CBS 6092 and CNRMA12.1409, which are differentiated by a mere 255 SNPs, exhibited markedly divergent *Dici2* contents (0.23% *versus* 2.09%). *Dici2* was identified as the most prevalent element in clades #2, #4, and #6. A notable exception was observed in strain CBS 564, which demonstrated a high level of *Dici2* (2.58%) in comparison to other strains of clade #6 (average: 1.39%). The presence of *Dici1* was only barely detected in strains of clade #4. Interestingly, the mosaic strains of clade #5 contained balanced copies of *Dici1* and *Dici2*.

**Fig. 4.**
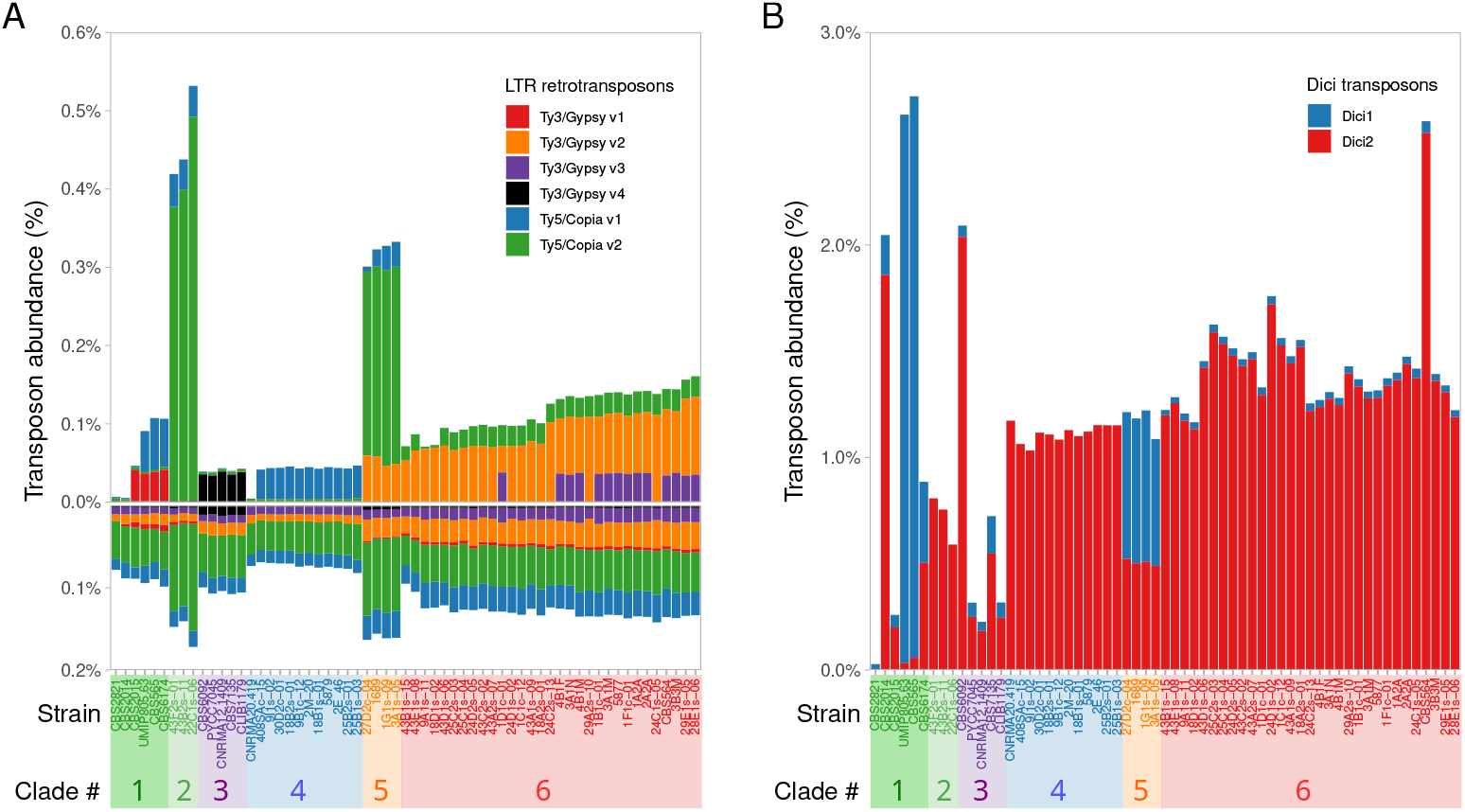
Transposon content in the genomes of the 61 *D. catenulata* strains. (A) abundance of LTR-retrotransposons (*Ty5* /*copia* and *Ty3* /*gypsy*), with the upper plot showing the prevalence of the *Ty* elements and the lower plot showing the prevalence of their corresponding LTRs; (B) abundance of *Dici1* and *Dici2* transposons. In both plots, the strains are grouped by clade in the same way

The structural features of *Dici1* and *Dici2* – including their similar size (≈ 4 kb), nearly identical TIR sequences (identical for the first 8 bp), and three CDSs in conserved order and orientation – indicate a common origin. However, the high nucleotide divergence between the two elements (52.2% identity) suggests that they have evolved over a long period. The remarkable sequence conservation of each element within and between strains points to recent bursts of amplification, as insufficient time has elapsed for mutations to accumulate between copies. Together, these observations indicate that *Dici* elements are biologically active and capable of rapid, strain-specific propagation in *D. catenulata* genomes. This hypothesis is further supported by RNA-seq data, which revealed that the three genes of *Dici* were expressed under our tested condition (Supplementary Fig. 2).

### 2.5 *Dici* belong to the *IS630-Tc1-mariner* group of transposons

The smallest and leftmost CDS of *Dici* (CDS1) encodes a protein comprising 143 or 146 (aa) in *Dici1* and *Dici2*, respectively. No function could be deduced from the detection of protein domains or sites using InterPro (Supplementary Table 6). For *Dici1*, the protein appears to contain a signal peptide that facilitates the transport of proteins across cellular membranes. Furthermore, no homologies were identified in protein databases (NCBI database: ClusterNR).

In the center of the *Dici* elements, CDS2 encodes for proteins of 477 (*Dici1*) and 473 aa (*Dici2*). The search in protein databases revealed similarities with uncharacterized fungal proteins containing an *IS630*-transposase domain (domain NF033545 in the Conserved Domain Database). A DDE-catalytic domain (InterPro entry IPR038717, Tc1-like transposase, DDE domain) was detected (Supplementary Table 6). This domain has been identified in the transposases of DD(E/D)-transposons, a group that belongs to Class II DNA transposons [2]. This group of TEs, which encompasses numerous superfamilies, is characterized by the occurrence of TIRs and TSDs, exhibiting variability in both their length and sequence [1]. Yuan and Wessler [41] proposed a revised classification system for DD(E/D)-transposons by defining a superfamily-specific “signature” consisting of conserved residues within the catalytic DD(E/D) domain. The protein sequence of CDS2 was found to match to the signature of the transposases of the *IS630-Tc1-mariner* superfamily transposons. This superfamily is ubiquitous in eukaryotes and fungi, and it has a simple structure consisting of two TIRs of variable length and one CDS encoding a transposase with a DD(E/D)-catalytic domain. Transposons belonging to this superfamily have a marked preference for insertions at adjacent TA dinucleotides [42], as was observed for *Dici* elements. Furthermore, a search in InterPro with the protein of CDS2 also detected a correspondence with homeodomain (HD)-containing proteins (InterPro entry IPR009057). This family of proteins contains a domain that binds DNA through a helix-turn-helix (HTH) structure found in a significant number of proteins, including transcription factors and transposases.

Structural analysis of the CDS2 sequence revealed characteristic features found in *IS630-Tc1-mariner* transposases [42]. Firstly, there was a pair of HTH motifs (PAIRED domain), each composed of three *α*-helices that were found in the N-terminal part of the protein (Fig. 2; Supplementary Fig. 3). This domain was reported to mediate the specific binding of the transposase to TIRs. Secondly, a GRPR-like motif was observed between the PAI and RED subdomains (GRTA in *Dici1* and GPKT in *Dici2*). Four putative nuclear localization signals (NLS), composed of basic amino acids (K or R), were predicted by PSORT. Finally, a DD34E catalytic domain, with 34 aa between the second aspartic acid (D) and glutamic acid (E), was localized in the C-terminal part of the protein. It is responsible of the catalytic activity required for the “cut-and-paste” mechanism of transpositions of the DNA transposons.

Finally, a search of protein databases using the sequence of CDS3 (347–398 aa) revealed similarity to uncharacterized fungal proteins annotated as containing an *IS630* transposase domain. A search in InterPro further indicated correspondence with homeodomain-containing proteins (Supplementary Table 6). These observations initially suggested that CDS3 might encode a transposase. However, a multiple-sequence alignment with CDS2 and related transposases revealed that the DDE catalytic triad was not conserved, indicating that CDS3 is likely a relic of a transposase gene (Supplementary Fig. 3). This finding points to the evolutionary history of *Dici* elements, suggesting that CDS3 represents a degenerate or inactivated copy that may have contributed to past transpositional activity within *D. catenulata* genomes.

In summary, the presence of three distinct CDSs contrasts with the typically compact architecture of *IS630-Tc1-mariner* elements, which have a single CDS that encodes the transposase.

### 2.6 *Dici*, a new family of *IS630-Tc1-mariner* transposons

A systematic search was performed against all available prokaryotic and eukaryotic genomes using the protein sequences of the *Dici1* and *Dici2* transposases as queries (see methods). A total of 22 putative homologues were identified in the kingdom of Fungi (Supplementary Table 8). Of these, 3 were detected in species belonging to the Ascomycota, which is the largest phylum of Fungi, while 19 were detected in the fungal phylum Mucoromycota.

The mean size of these fungal elements was found to be 1,749 base pairs, with the exception of *Dici1* (4,031 bp), *Dici2* (4,165 bp), and the transposon detected in *Mucor piriformis* (FUN-MUPI-2), which has an unusual size of 6,758 bp (Supplementary Fig. 4; Supplementary Table 8). This later transposon exhibited a complex structure, characterized by two inverted-repeat regions of 1,627 bp at both extremities, which harbor two identical transposase CDSs (438 aa). The TIR sequences displayed a considerable degree of size variability, ranging from 8 to 187 base pairs. The most prevalent TIR sequences were found to initiate with CAGTA (*n* = 9) or CACCT (*n* = 7). A characteristic dinucleotide TA duplication was invariably present at each extremity, a common characteristic of transposons of the *IS630-Tc1-mariner* group.

Their corresponding transposases are generally encoded by a single CDS ranging from 432 to 478 aa. The characteristic domains previously observed in the *Dici* transposases were conserved, namely the DNA-binding (PAIRED) and DD34E catalytic domains (Supplementary Fig. 3). The four putative NLS patterns were found to be conserved. Notably, the rightmost pattern (KKRK), located in the core of the DD34E catalytic domain, was detected in all transposases. The GRPR-like motif, located between the PAI and RED subdomains, was identified in multiple forms, with GRP(P/A) being observed in 14 transposase sequences. Finally, The mean pairwise sequence identity between the transposase sequences is 40%, rising to 52% when the analysis was restricted to the catalytic domain (Fig. 5).

**Fig. 5.**
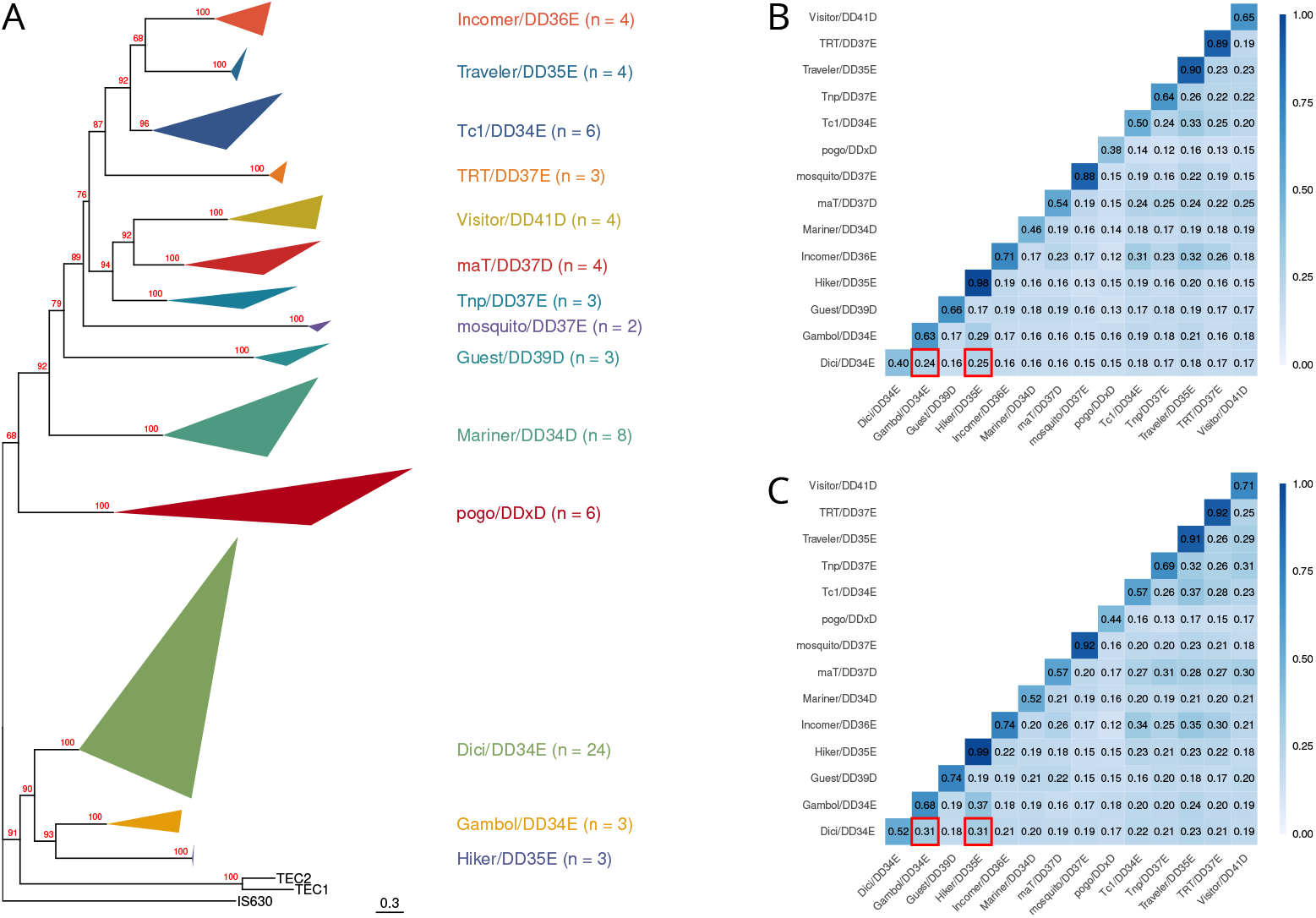
Comparison of the *Dici* transposases with transposases belonging to the *IS630-Tc1-mariner* superfamily. (A) phylogenetic tree generated using the full-length transposase sequences and the maximum-likelihood method with a bootstrap approach (bootstrap values are shown in red). The transposase of *IS630* was used as an outgroup. A version of this tree, with expanded clades and complete transposase names, can be seen in Supplementary Fig. 5; (B and C) heatmaps showing pairwise protein identities between the transposases of different families. The average off all pairwise identities measured between two transposon families is indicated by numbers in the cells. Comparison was measured using either (B) the full-length transposases, or (C) restricted to the catalytic domain. The values obtained for the comparisons of the transposases of the Dici family with those of the Gambol and Hiker are highlighted by a red square. Refer to the Supplementary Table 8 for a detailed description of the transposase name and family

A phylogenetic approach was used to classify the *Dici* transposases alongside representatives of the *IS630-Tc1-mariner* group of transposons (Fig. 5). The results revealed that *Dici* transposases formed a distinct clade, with 100% bootstrap support, within the Gambol/DD34E [43] and Hiker/DD35E [44] transposase families. These transposase families were found to be distinct from the other known transposon families in the *IS630-Tc1-mariner* group [44]. Pairwise sequence comparisons between the different families confirmed this relationship with the Gambol and Hiker transposases (Fig. 5). The highest conservation was observed when comparing the full-length *Dici* transposases with those of Gambol/DD34E (24%) and Hiker/DD35E (25%), rising to 31% when the analysis was restricted to their catalytic domains. In summary, these results suggest that *Dici* forms a new family, which is related but distinct from the Gambol and Hiker families.

## 3 Discussion

The present study has revealed two new transposable elements, named *Dici1* and *Dici2*, in the *D. catenulata* genomes. Along with newly discovered fungal transposons, they form a new family, named *Dici*, within the highly diverse *IS630-Tc1-mariner* group of DNA transposons [40]. The *Dici* family has, to date, only been observed within the fungal kingdom. According to phylogenetic analysis of transposases, this family forms a monophyletic clade close to the Gambol/DD34E and Hiker/DD35E elements, which were previously considered distinct from the main Tc1/DD34E family [43].

Transposons are commonly associated with genome rearrangements and unique chromosome features [12]. We detected extensive large-scale chromosomal rearrangements with no more collinearity between chromosomes of the different strains. Of particular significance is the observation that many chromosomal breakpoints coincide with the insertion sites of *Dici* elements, suggesting that transposition activity may have contributed significantly to the observed genomic instability. Comparable patterns have been documented in the *Saccharomyces* yeasts, where the rearrangements appeared to result from ectopic recombination between LTR-retrotransposons (*Ty* elements) or other repeated sequences [45].

The pronounced genetic structure observed for *D. catenulata* reveals distinct clades representing deeply diverged lineages with limited gene flow. This pronounced differentiation indicates that historical barriers to genetic exchange have persisted over extended evolutionary timescales. Here, the substantial chromosomal rearrangements that generated divergent karyotypes likely played a role by disrupting meiotic pairing, acting as intrinsic barriers to recombination and preventing gene flow among the *D. catenulata* population. Such chromosomal rearrangements have been proposed to promote hybrid infertility [46] and to lock in adaptive alleles [47], thereby accelerating lineage divergence.

The current distribution and high sequence conservation of *Dici* elements across *D. catenulata* clades may be explained by two main scenarios. In the first scenario, both *Dici1* and *Dici2* transposons were acquired by a common ancestor prior to clade diversification, as suggested by their presence in all clades. A subsequent burst of *Dici* transposition may have contributed to early genomic rearrangements and potentially triggered lineage divergence. The absence of genetic recombination between rearranged genomes likely promoted the accumulation of mutations within each clade. The observed variation in *Dici* copy number among strains within the same clade further supports the occurrence of subsequent strain-specific transposition dynamics. In the second scenario, the high nucleotide conservation of *Dici* elements both within and between clades is consistent with a more recent acquisition event, which was subsequently followed by a rapid burst of transposition. The detection of both *Dici1* and *Dici2* elements across all lineages may be attributed to multiple independent introductions or a single acquisition event that was subsequently disseminated through horizontal transfer among lineages. This hypothesis is further supported by the observation that some strains, isolated from the same PDO cheeses, were found in different clades, a pattern that may have facilitated such horizontal exchanges. In this scenario, the *Dici* elements likely invaded already differentiated genomic backgrounds and later expanded, contributing to the structural variations observed today.

Taken together, our findings establish *D. catenulata* as a powerful model for dissecting the mechanisms of DNA transposon proliferation and elucidating how mobile genetic elements drive genome plasticity. These processes may be pivotal in fungal evolution, by facilitating rapid adaptation and promoting speciation. Beyond providing fundamental insights, these results may provide key information for investigating the emergence of opportunistic fungi in both clinical and food safety contexts.

## 4 Methods

### 4.1 Strains and culture media

The 61 *D. catenulata* strains used in this study were obtained from international collections (CLIB, CBS and PYCC strains), or isolated from cheeses as part of the MetaPDOcheese project [22]. Other were kindly provided from their own collections by Françoise Irlinger from INRAE SayFood (Palaiseau, France), Marie Desnos-Ollivier from Pasteur Institute (Paris, France), and Nicole Desbois-Nogard from the University Hospital of Martinique (Fort-de-France, France) (Supplementary Table 1). Yeast strains were maintained on YPD medium (10 g/L yeast extract, 20 g/L bacto peptone, 20 g/L glucose, 20 g/L agar).

### 4.2 Isolation and identification of yeast strains from cheese

In a previous study, 2,316 subsamples of rinds and cores from 1,158 cheeses of 44 French PDO cheeses were collected and the diversity of their microbial community was assessed using a metabarcoding approach [22]. The PDO cheeses were anonymized and randomly coded from PDO #1 to PDO #44. Based on the metabar-coding results, we selected PDO cheeses in which *D. catenulata* was present with an abundance greater than 10%, which represents 5.5% of the 2,316 cheese samples. An aliquot of each cheese was plated on YPD and Wallerstein (Thermo Fisher Scientific) media containing 100 µg/mL chloramphenicol and incubated 48 h at 28 °C. Individual colonies were identified taxonomically by sequencing the D1/D2 and ITS regions using primers ITS1 (5’-TCCGTAGGTGAACCTGCGC-3’) and NL4 (5’-GGTCCGTGTTTCAAGACGG-3’) as previously described [48]. In total, 35 isolates of *D. catenulata* were selected from 12 PDO cheeses, corresponding to 28 different producers. Seven producers provided us with two seasonal cheeses (18B, 24C, 24D, 25B, 25C, 43A and 43B). Only one strain per cheese was analyzed in this study.

### 4.3 Flow cytometry

Yeast cells were prepared for flow cytometry analysis as previously described [48]. DNA labeled with SYTOX R green (Invitrogen) was quantified on a C6 Accuri (Ann Arbor, MI, United States) flow cytometer as previously described [49], with an excitation wavelength of 488 nm and an emission wavelength of 530 nm. Acquisition was performed on 30,000 events observed with gating on forward scatter/side scatter signals. The flow rate was set to about 2,000 events per second (medium flow, 35 mL/min; core, 16 mm).

### 4.4 Pulsed-field gel electrophoresis

Yeast karyotyping was achieved by contour-clamped homogeneous electric field (CHEF) gel electrophoresis. Plugs of yeast chromosomes were prepared as described elsewhere [50]. The CHEF-DR III apparatus (Bio-Rad, Hercules, CA, United States) was set to 5.2 V/cm with pulses of 90-120 s for 12 h and then to 4 V/cm with pulses of 120-360 s for 24 h, and samples were run on 1% SeaKem^®^ Gold Agarose gels (Lonza, Rockland, ME, USA) in 0.5 x TAE buffer at 12 °C. The chromosomes of *Saccharomyces cerevisiae* CLIB 112 (= YNN295) were used as size markers. The agarose gels were stained with ethidium bromide (0.5 mg/mL) and washed with water before visualization under UV.

### 4.5 DNA extraction

Yeasts were cultured in 10 mL YPD medium at 28 °C with shaking at 220 rpm for at least 36 h [49]. Genomic DNA used for sequencing with Illumina technology was extracted based on an in-house protocol involving mechanical (vortexing for 4 min with 0.3 g glass beads) and chemical lysis (Tris-HCl 10 mM pH8, EDTA 1 mM, NaCl 100 mM, Triton 2% and SDS 1%), as previously described [48]. To eliminate small fragments from the genomic DNA, a purification was performed using AMPure XP beads (Beckman Coulter Genomics, Danvers, MA, USA).

### 4.6 Illumina library preparation and sequencing

Libraries were prepared with NEBNext DNA Modules Products (New England Biolabs, Ipswich, MA, USA) and an “on beads” protocol developed by Genoscope. Genomic DNA (250 ng) was used as starting material. After adapter ligation, the ligated product was amplified by 12 cycles of PCR with the Kapa Hifi Hotstart NGS library Amplification kit (Roche, Basel, Switzerland), followed by purification with AMPure XP beads (Beckman). Libraries were sequenced with an Illumina NovaSeq 6000 instrument (Illumina, San Diego, CA, USA), in paired-end mode, generating 150-base reads. Illumina short-read sequencing statistics are reported on Supplementary Table 9. Average sequencing depth across the 61 *D. catenulata* strains is 140X.

Following Illumina sequencing, we applied an in-house quality control process to the reads that passed the Illumina quality filters. The first step discarded low-quality nucleotides (*Q <* 20) from both ends of the reads. The Illumina sequencing adapter and primer sequences were then removed from the reads, and any reads shorter than 30 nucleotides after trimming were discarded. These trimming and removal steps were achieved using in-house software based on the FastX package (https://github.com/institut-de-genomique/fastxtend). In the final step, read pairs that mapped to the phage *ϕX* genome were discarded using the SOAP aligner [51], along with the reference genome of the *ϕX*174 enterobacterial phage (GenBank: NC_001422.1). Short Illumina reads were subjected to post-processing to remove low-quality data, as previously described [52, 53].

### 4.7 SNP calling

The Illumina reads for each *D. catenulata* strain were mapped using bwa-mem2 version 2.0pre1 [54] to the genome obtained for the type strain CBS 565^T^. The large segmental duplication (388 kb) observed in chromosome F (DICA0_F:3198109-3586457) was masked to prevent null mapping quality being obtained in duplicated regions, which would impact subsequent variant detection.

We used Genome Analysis Toolkit (GATK) v4.5.0.0 for variant calling (Haplotype-Caller parameters: `--sample-ploidy 2, --emit-ref-confidence GVCF`), with hard filtering according to GATK best practice [55]. The genotyping pipeline produced a VCF file of 508,773 variants, which included 436,405 SNPs and 72,368 indels identified in the 61 yeast samples.

Only SNPs were retained for phylogenetic analysis and population genomics. Those with a missing genotype above 10% or a minor allele frequency below 0.03 were removed. Genotypes with quality and read depths below 30 were masked. SNPs were filtered using vcftools version v0.1.16 [56], using the following parameters: `--minGQ 30 --minDP 30 --maf 0.03 --max-missing 0.9`. The resulting filtered dataset contained 357,967 SNPs. Initial analysis revealed the presence of heterozygous SNPs even in haploid genomes (on average 1,150 positions per genome). These SNPs, which were mostly clustered, were removed if they fell within a region containing more than seven heterozygous SNPs within a 1,000 bp window. The final dataset contained 351,942 SNPs.

### 4.8 Phylogenetic analysis

For the purpose of phylogenetic analysis, the VCF SNP file (351,942 SNPs) was used to generate sequences in FASTA format, with IUPAC ambiguity codes (Y, R, W, S, K, M) used for heterozygous positions when necessary. Subsequently, a phylogenetic tree was generated using IQ-TREE v2.3.1 (parameters: `-T AUTO -m MFP`), with a maximum-likelihood approach [57]. The best-fit nucleotide substitution model was automatically selected by ModelFinder [58]. Branch support was assessed using an ultra-rapid bootstrap approximation (parameter `-B 1000`).

### 4.9 Population genomics

Population structure inference was performed using fastStructure [59]. The VCF SNP file (351,942 SNPs) was converted to PED format using PLINK v1.90b7.2 [60], filtering positions that are in linkage disequilibrium (*LD*_1*/*2_ *>* 0.5, PLINK parameters: `--indep-pairwise 50 5 0.5`). This resulted in a set of 34,101 SNPs. Twenty runs of fastStructure were performed with the prior set to “simple” and summarized using CLUMPP v1.1.2 [61]. Plotting was carried out using R v4.5.0 [62].

The AMOVA method [63] implemented in the R package poppr v2.9.7 [64] was used for the detection of population differentiation. Permutation tests, which involve the random reallocation of individuals or haplotypes to different groups, were performed using the randtest function from the ade4 v1.7-23 package [65].

Population genetics statistics (*π, d*_*XY*_, *F*_*ST*_ and Tajima’s *D*) were computed on the VCF SNP file (351,942 SNPs) using the R package PopGenome v2.7.7 [66] and a window of 10,000 bp. The values were averaged over the windows to obtain genome-wide values.

### 4.10 Transposon content

For each strain, the transposon content was estimated by calculating the ratio of reads mapped to each element, divided by the total number of reads mapped to the genome. This was achieved by mapping short reads for each of the 61 yeasts using bwa-mem2, and by using a FASTA file containing one copy of each transposon family (see Results). Read counts for each element were obtained using featureCounts v2.0.6 in the Subread package [67]. The number of reads that mapped to the genome was obtained using samtools v.1.15.1 (samtools view parameters: `-c -G 13 -F 256`) with the BAM files generated by bwa-mem2 (Supplementary Table 9).

### 4.11 Chromosomal rearrangements analysis

SynChro was used to reconstruct synteny blocks from pairwise comparisons of the five annotated genomes [68]. MUMmer4 package was used to generate global alignments of the genomes [69]. The alignments were further filtered using the delta-filter utility (MUMmer4) with parameters `-l 5000` (exclude regions ≤ 5 kb) and `-m`. Alignment coordinates were extracted with the show-coords utility to generate a tabular file. Circular representation of chromosomal rearrangements was performed in R using the package circlize v0.4.16 [70].

### 4.12 *Dici* transposons characterization

Functional analysis of the three protein sequences of *Dici* elements was performed using InterPro (https://www.ebi.ac.uk/interpro). Protein secondary structures predictions were performed using the PSIPRED version 4 (http://bioinf.cs.ucl.ac.uk/psipred). Nuclear localization signal (NLS) predictions were performed using the PSORT II (http://psort.hgc.jp).

### 4.13 Mining of *Dici* transposons

To assess the taxonomic distribution of *Dici* transposons in genomes, *Dici1* (DICA0_E02476) and *Dici2* (DICA4_A06172) transposase sequences were used as a query to search the National Center for Biotechnology Information (NCBI) non-redundant (nt) and eukaryotic reference genomes (Eukaryote Refseq Genomes) nucleotide databases (downloaded on 23/05/2025) using TBLASTN (version 2.15.0) with an E-value cutoff of 10^−10^. The reported sequences were kept only if a catalytic DDE domain was identified using InterProScan version 5.71-102.0 (parameters: `--applications PFAM`, search domain: PF13358).

To determine the boundaries of these elements, the best hits sequences were recovered with 2-kb flanking regions, aligned using MAFTT v7.453 [71] with parameters `--maxiterate 1000 --genafpair`. Transposon boundaries were then checked manually for the presence of terminal inverted repeats (TIR) and target site duplications (TSD). The resulting transposon sequence was searched against its host genome using BLASTN (E-value cutoff 10^−40^), to estimate copy number. All hits obtained that were *>* 40% in size and had *>* 80% identity were used to calculate the copy number as proposed by [1].

### 4.14 *Dici* transposases phylogeny

Multiple sequence alignments of transposase sequences (full-length or catalytic domain) were obtained using MAFFT (parameters: `--maxiterate 1000 --globalpair`). ClipKIT version 1.3.0 [72] was used to remove gaps and keep only phylogenetically informative sites in the alignment (parameter: `-m kpi-smart-gap`). The phylogenetic trees were inferred using the maximum-likelihood method within the IQ-TREE version 2.3.1 [57]. The best-suited protein substitution model was selected using ModelFinder [58] embedded in IQ-TREE (parameters: `-T 10 -mset LG,JTT,WAG,Dayhoff,NQ.yeast,Q.yeast`). The reliability of the maximum likelihood trees was estimated using the ultrafast bootstrap approach with 1,000 replicates (parameter: `-B 1000`).

Family-wise transposase comparisons were obtained using the multiple alignments (full-length or catalytic domain) generated with MAFFT. The alignment files were imported and the matrices of pairwise identity were obtained using the seqinr R package (v4.2-36) [73]. Then, identity values for each pair of transposon families were averaged.

## Supporting information

Supplementary results, materials, procedures and figures

## Supplementary information

### Supplementary text and figures

Additional results, materials, procedures and figures. (PDF)

- **Supplementary Figure 1**. Chromosomal rearrangements in the *D. catenulata* genome.
- **Supplementary Figure 2**. Expression of the three *Dici* transposon genes was revealed by RNA-seq data.
- **Supplementary Figure 3**. Complete alignments of the *Dici* transposases.
- **Supplementary Figure 4**. Characteristics the transposon detected in *Mucor piriformis*.
- **Supplementary Figure 5**. Phylogenetic tree of the *IS630-Tc1-mariner* transposases with clades not collapsed.
- **Supplementary Figure 6**. Structure of the mating-type like (MTL) locus of *D. catenulata*.

### Supplementary tables

Additional spreadsheet documents (XLSX)

- **Supplementary Table 1**. *D. catenulata* strains used in the study. Also includes accession numbers for sequencing and assembled genome data.
- **Supplementary Table 2**. Genetic diversity and differentiation statistics measured in *D. catenulata*.
- **Supplementary Table 3**. Assembly and annotation statistics of *D. catenulata* assembled genomes.
- **Supplementary Table 4**. Genes associated with drug resistance and virulence detected in *D. catenulata*.
- **Supplementary Table 5**. Characteristics and chromosomal positions of the RNA transposons discovered in *D. catenulata* genomes.
- **Supplementary Table 6**. Characteristics and chromosomal positions of the *Dici* transposons discovered in *D. catenulata* genomes.
- **Supplementary Table 7**. List of synteny blocks detected between the *D. catenulata* CBS 565 genome and those of the other four cheese strains.
- **Supplementary Table 8**. Characteristics of the *Dici* transposons newly identified in Fungi. Also contains the list of the *IS630-Tc1-mariner* transposase sequences used to generate the phylogenetic tree.
- **Supplementary Table 9**. Sequencing statistics for the five *D. catenulata* assembled genomes. Also contains short-reads mapping statistics obtained for the 61 *D. catenulata* stains.

## Acknowledgements

The authors would like to thank the following people for providing strains: Françoise Irlinger (INRAE SayFood, Palaiseau, France); Marie Desnos-Ollivier (Pasteur Institute, Paris, France); Nicole Desbois-Nogard (University Hospital Center of Martinique Island, Fort-de-France, France); CIRM-Levures (https://cirm-levures.bio-aware.com); and the USDA-ARS Culture Collection (NRRL). We would also like to thank Hugo Devillers (INRAE, Montpellier, France) for his technical assistance with bioinformatics.

## Author contribution

Conceptualization: CN; Data curation: FB, CN; Formal analysis: FB, CN, MW, JMA; Investigation: MP; Methodology: CC, MW, JMA; Project administration: CN, JMA; Resources: MP, CN; Supervision: CN, JMA; Visualization: FB, CN; Writing – original draft: FB, CN; Writing – review & editing: FB, CN, JMA, MW, CC.

