## Supplementary results, materials, procedures and figures for "*Dici*: a novel DNA transposon reshaping the genome of the opportunistic yeast *Diutina catenulata*"

#### Supplementary Information

Frédéric Bigey

November 6, 2025

**3 Supplementary figures** **7**

**4 Supplementary tables** **14**

**References** **15**

### 1 Supplementary results

#### 1.1 Preparation of *D. catenulata* yeast collection

New *D. catenulata* isolates were collected from twelve french Protected Designation of Origin (PDO) cheeses (PDO #1, #3, #9, #18, #22, #24, #25, #27, #28, #29, #30 and #43) produced by twenty-eight different producers, during a previous project [1]. A selection of isolates was made from cheeses produced by the same producer in two separate seasons (producers 18B, 24C, 24D, 25B, 25C, 43A and 43B). This resulted in a collection of 35 *D. catenulata* isolates. Additionally, we supplemented this set with isolates from international yeast collections and laboratories. These were selected from eight geographical and five ecological origins, primarily animal/human ($n = 9$) or dairy ($n = 14$). The resulting yeast collection comprises 61 *D. catenulata* strains (Supplementary Table 1).

#### 1.2 New genomic resources for *D. catenulata*

Five haploid *D. catenulata* strains (CBS 565$^{T}$, 43B2s-11, CLIB 1179, 25B2s-01 and 1F1c-01) were selected for genome sequencing and assembly. These strains are representative of the clades identified during the phylogenetic analysis (see main text). A hybrid sequencing strategy was employed, in which short reads were coupled with long reads (see methods section below). The *D. catenulata* assembled nuclear genome ranges in size from 14.0 Mb ($n = 4$) to 14.7 Mb ($n = 1$; CBS 565) and comprises six near-complete chromosomes ranging in size from 0.8-1.2 Mb for the shortest, to 3.5-4.2 Mb for the largest (Supplementary Table 3), which is compatible with the sizes observed in the karyotypes (see manuscript). Telomere repeats of 26 bp were identified at the extremities of some contigs, thereby confirming that at least two telomere-to-telomere chromosomes were obtained for each strain. The rDNA loci were located at three contig extremities that lacked telomere repeats. The 53.5% GC-content nuclear genomes of *D. catenulata* encode approximately 6,300 CDSs. The BUSCO score was found to be 98.1% on average after the annotation was manually curated. In addition to protein-coding genes, approximately 300 tRNA genes were predicted. Summary statistics are presented in Supplementary Table 3.

#### 1.3 Analysis of the mating-type like locus

Synteny at the MTL (mating-type like) locus of *D. catenulata* (Supplementary Fig. 6) was conserved with that described for *Candida anglica* [2] and *C. railenensis* [3], between *RCY1* and *PIK1* genes. In comparison to *C. albi-*

*cans* and *C. parapsilosis*, the CDS corresponding to CAALFM_C501730WA or CPAR2_303610 was not observed [4]. The transcription activator MTL$\alpha$1 gene was observed in *D. catenulata* CBS 565 and 1F1c-01, while no evidence of either MTLa or MTL$\alpha$ genes was found in strains 43B2s-11, CLIB 1179, and 25B2s-01. The MTL$\alpha$2 gene was identified outside the MTL locus. Taken together, these findings suggest that all, or at least the majority, of *D. catenulata* isolates possess a defective mating pathway.

#### 1.4 Analysis of the mitochondrial genomes

Complete circular mitochondrial genomes (Supplementary Table 3), ranging from 20,934 to 22,153 bp (average GC content is 25.8%), were identified in *D. catenulata* strains CBS 565 (accession: OZ261072), 43B2s-11 (accession: OZ261100) and 25B2s-01 (accession: OZ262066). The annotation revealed 15 CDS and 25 tRNA, all of which are in the same orientation. A putative sequence of an intron-encoded DNA homing endonuclease with a LAGLIDADG domain was found in the gene coding NADH dehydrogenase subunit 5 (nad5) in strains 43B2s-11 and 25B2s-01 only.

#### 1.5 Antifungal resistance and virulence genes were detected in *D. catenulata*

A search of genes, associated with antifungal resistance and virulence in the pathogenic yeast *C. albicans*, was conducted (Supplementary Table 4). Both major antifungal drug targets, ERG11 (lanosterol 14-$\alpha$-demethylase) for azoles and FKS1 ($\beta$-1,3-glucan synthase) for echinocandins, are encoded by single-copy genes. In contrast, the genes coding for drug efflux transporters, which confer resistance to numerous chemicals, including azoles, showed notable amplification, with *CDR1* presents in three copies across all strains and *MDR1* occurring in three to four copies. The transcription factors that regulate their expression were detected as single copies for *TAC1*, and with a limited variation for *MRR1* (0–2 copies). Other genes coding for transcription factors and regulators involved in morphogenesis (*CPH1*, *EFG1*) and filamentous growth (*RBF1*, *TUP1*), which have critical biological functions for fungal virulence and pathogenicity, were detected mainly in single copies. Several genes coding for hydrolytic enzymes associated with pathogenicity showed expansion. *PLB1*, coding for phospholipase B1, involved in host membrane degradation, was present in three copies in all genomes. Members of the SAP family (secreted proteinases that degrade host proteins) showed gene copy number variation, with *SAP5* ranging from three to five copies.

It is noteworthy that the distribution of these drug resistance and virulence-associated genes did not differ between the clinical isolate CBS 565 and the cheese-derived strains. This suggests that the genetic potential for antifungal resistance and pathogenicity is conserved in environmental and clinical isolates. Overall, this pattern indicates that while the regulatory backbone of resistance and virulence remains mainly conserved as a single copy, the amplification of effector genes, particularly multidrug transporters and hydrolytic enzymes, may contribute to the adaptive potential of *D. catenulata*.

### 2 Supplementary materials and procedures

#### 2.1 DNA and RNA extraction

To extract yeast genomic DNA for long-read sequencing, strains were cultured in 10 mL YPD medium at 28 °C with shaking at 220 rpm for at least 36 h [2]. The extraction method was adapted from a protocol developed at Genoscope [5]. Briefly, yeast cells are lysed with zymolyase to generate spheroplasts, which are then chemically lysed in the presence of SDS. Proteins are then removed by precipitation with potassium acetate. Finally, the DNA is precipitated with isopropanol, washed with ethanol and resuspended in TE (10 mM Tris-HCl, 1 mM EDTA).

To extract total RNA, strains were cultured overnight in 10 mL YPD medium at 28 °C with shaking (220 rpm). Cell concentration was then estimated by spectrophotometry and $10^9$ cells were harvested. The TRIzol method [6] was used to isolate total RNA as previously described [7]. Briefly, cells were quickly washed with 750 mL cooled (4 °C) DEPC-treated water, pelleted, frozen in a -80 °C methanol bath and mechanically lysed through vortexing with glass beads in 400 µL TRIzol™ (Life Technologies) at 4 °C for 15 min. The liquid phase was collected and TRIzol added to a 4 mL final volume, with 800 µL chloroform. The mixture was then vortexed and centrifuged (9,000 g for 15 min). The supernatant was centrifuged again (2,000 g for 2 min). RNAs were precipitated with 2 mL cooled isopropanol and incubated for 10 min at -20 °C before centrifugation (9,000 g for 10 min). The resulting pellet was washed twice with 75% ethanol and then dissolved in 150 µL of nuclease-free water (Qiagen). Total RNA from 100 µg aliquots was purified with a RNeasy mini kit (Qiagen). RNAs were eluted with 2 × 30 µL of the provided RNAse-free water.

#### 2.2 Long reads library preparation and sequencing

Genomic DNA (9 µg) was purified from five strains of *D. catenulata* (CBS 565, 43B2s-11, CLIB 1179, 25B2s-01, 1F1c-01) using the Short Read Eliminator XL

Kit (Pacific Biosciences, Menlo Park, CA, USA). A library was prepared with 2 µg purified genomic DNA as the starting material, following the "1D Native barcoding genomic DNA (with EXP-NBD104 and SQK-LSK109)" protocol provided by Oxford Nanopore Technologies (Oxford Nanopore Technologies Ltd, Oxford, UK). The sample (pooled with five other barcoded samples) was sequenced using a R9.4.1 MinION flow cell. The reads were basecalled using Guppy version 4.2.5. ONT long-read sequencing statistics are reported on Supplementary Table 9.

#### 2.3 Illumina RNA-seq library preparation and sequencing

Libraries were prepared from 100 ng total RNA extracted from five strains of *D. catenulata* (CBS 565, 43B2s-11, CLIB 1179, 25B2s-01, 1F1c-01) using the NEBNext Ultra II Directional RNA Library Prep for Illumina (New England BioLabs) according to the manufacturer's protocol. The libraries were sequenced using an Illumina NovaSeq 6000 instrument (Illumina, San Diego, CA, USA) in paired-end mode, generating 150-bp reads.

#### 2.4 Long read-based genome assembly

The raw Nanopore reads for the five strains of *D. catenulata* were assembled using NECAT with default parameters [8] and a genome size of 20 Mb. The resulting output was polished once with Racon [9] using Nanopore reads, then once with Medaka (model r941_min_high_g360) using Nanopore reads, and twice with Hapo-G 1.3.4 [10] using Illumina short reads.

#### 2.5 Genome annotation and manual curation

Tandem repeats in the genome assemblies were masked using Tandem Repeat Finder [11], while simple repeats and known repeats included in RepBase were masked using RepeatMasker (`https://repeatmasker.org/`) with the parameter `-species Saccharomycetes` [12].

Gene prediction was performed using several proteomes downloaded from NCBI: including *Candida railenensis*, *Debaryomyces hansenii*, *Diutina rugosa*, *Huiozyma naganishii* (formerly *Kazachtania naganishii*) and *Kazachstania africana*, which were aligned against the genome in a two-step strategy. To rapidly locate the regions in the genome corresponding to these proteins, we used BLAT version 36 with default parameters [13]. We retained the best match and matches with a score $\geq$ 90% of the best match score. The alignments were then refined using Genewise version 2.2.0 with default parameters [14], as this approach is more accurate for detecting intronexon boundaries. We retained alignments if $> 75\%$ of the length of the protein could be aligned with the genome.

RNA-seq short reads were used to detect expressed and/or specific genes. The reads were mapped onto the genome assembly using HISAT2 version 2.2.1 with default parameters [15]. The resulting BAM file was sorted and Stringtie version 2.2.1 [16] was applied with the following parameters: `-p 16 -v -m 150`. Only the most highly expressed transcript at each genomic locus was retained.

The protein and RNA-seq alignments were combined using Gmove, an easy-to-use predictor that does not require precalibration [17] and is available on GitHub (https://github.com/institut-de-genomique/Gmove). Genes with $> 50\%$ untranslated regions and a coding sequence (CDS) length of less than 300 bp were excluded.

The RNA-seq data was used for expert manual curation of gene models. This allowed for identification of missing genes in the assembled genomes and correction of intron splice sites. To this end, RNA-seq reads were mapped against the genome assemblies using HISAT2 with the following parameters: `--rna-strandness FR --max-intronlen 3000`. The results of RNA-seq mapping (BAM files) were displayed using the Artemis genome browser v18.2.0 [18]. Manual changes were made to the structural annotations within the browser to correct intron splice sites. The completeness of the annotation was assessed using BUSCO v5.3.1 [19], with the the following parameters: `-m protein` and `saccharomycetes_odb10` lineage dataset as the reference (2,137 proteins). Functional annotation was performed using the in-house software go-FAnnoT (https://github.com/hdevillers/go-fannot), which assigns annotations based on homologies found in the curated UniProt database (release 2025_01).

In addition to predicting protein-coding genes, tRNA genes were identified using tRNAscan-SE v2.0.5 [20]. A BLASTN search was also conducted to detect the presence of a complete rDNA unit, including 18S, 5.8S, 26S and 5S rRNA genes. The reference sequences of *Candida anglica* [2] were used for this analysis.

#### 2.6 Sequencing and genomic data accessibility

The raw sequencing data and annotated genomes have been deposited in the European Nucleotide Archive (ENA) at EMBL-EBI under project accession number PRJEB94530. For each strain, the corresponding accession numbers are available in Supplementary Table 1.

#### 3 Supplementary figures

- **Supplementary Figure 1**
  Chromosomal rearrangements in the *D. catenulata* genome.
- **Supplementary Figure 2**
  Expression of the three *Dici* transposon genes was revealed by RNA-seq data.
- **Supplementary Figure 3**
  Complete alignments of the *Dici* transposases.
- **Supplementary Figure 4**
  Characteristics the transposon detected in *Mucor piriformis*.
- **Supplementary Figure 5**
  Phylogenetic tree of the *IS630-Tc1-mariner* transposases with uncollapsed clades.
- **Supplementary Figure 6**
  Structure of the mating-type like (MTL) locus of *D. catenulata*.

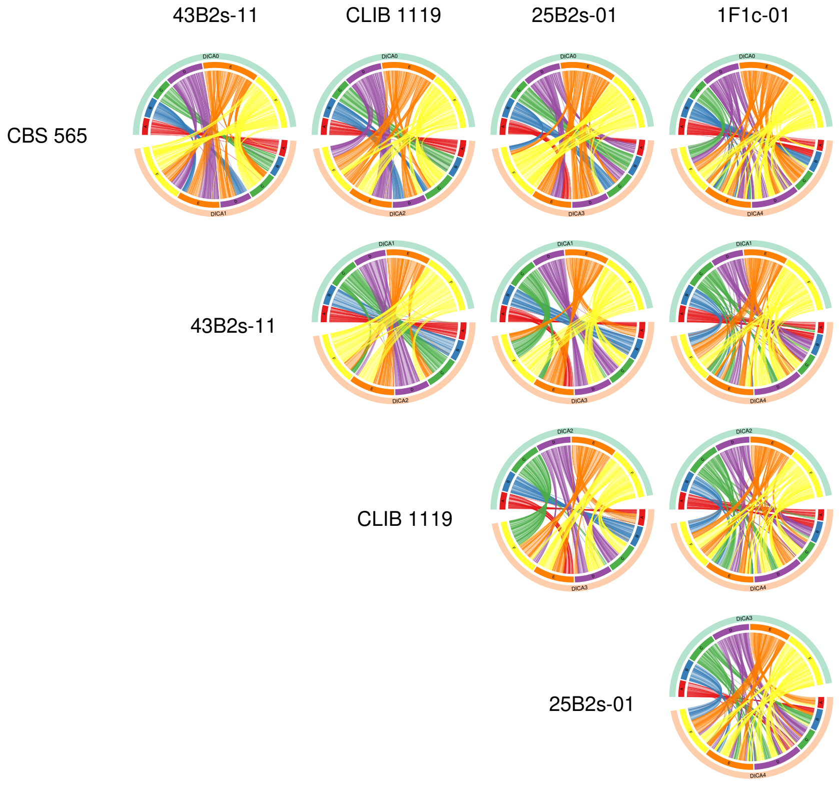

**Supplementary Figure 1.** Chromosomal rearrangements in the *D. catenulata* genome. Circular plots showing the pairwise genome alignments of the five *D. catenulata* strains. The lines between the genomes represent aligned regions exceeding 5 kb. Lines have the same colors as the chromosomes of the genome on the top of the circle, corresponding to the strain on the left

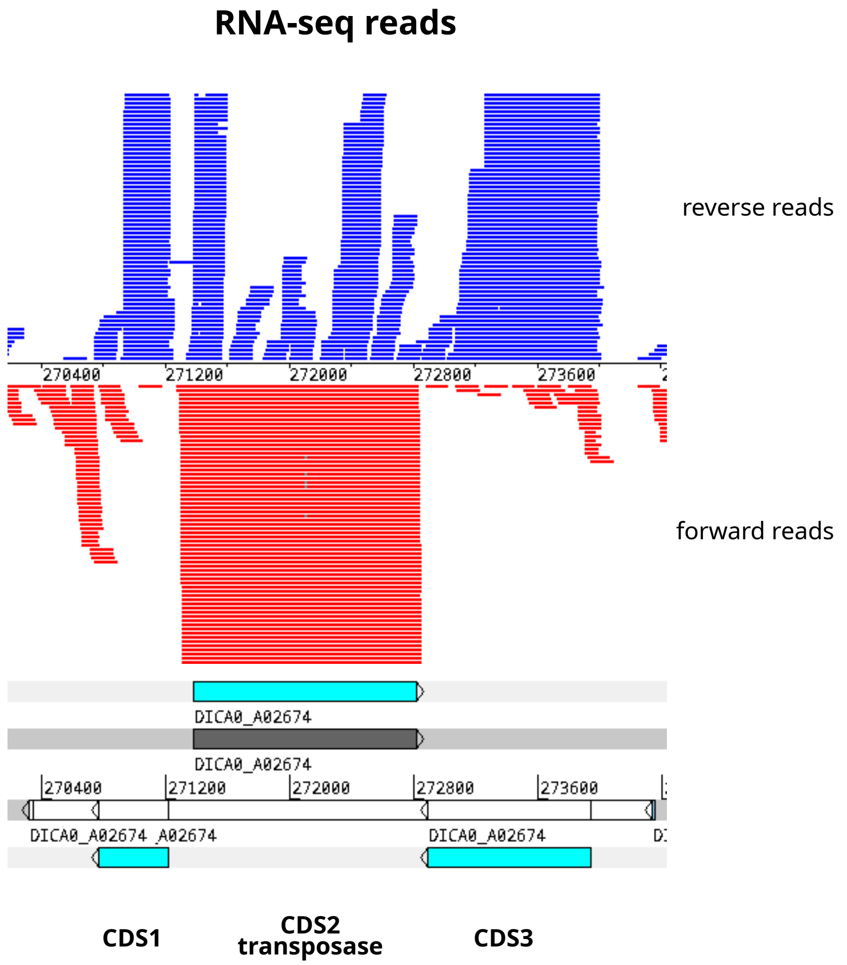

**Supplementary Figure 2.** Expression of the three *Dici* transposon genes was revealed by RNA-seq data. The RNA-seq reads obtained from the mRNA extracted from the *D. catenulata* strain CBS 565 were mapped to the CBS 565 genome. The mapped reads over the *Dici1* transposon DICA0_A02674 were visualized using the Artemis software program. A similar expression was observed for the *Dici2* genes.

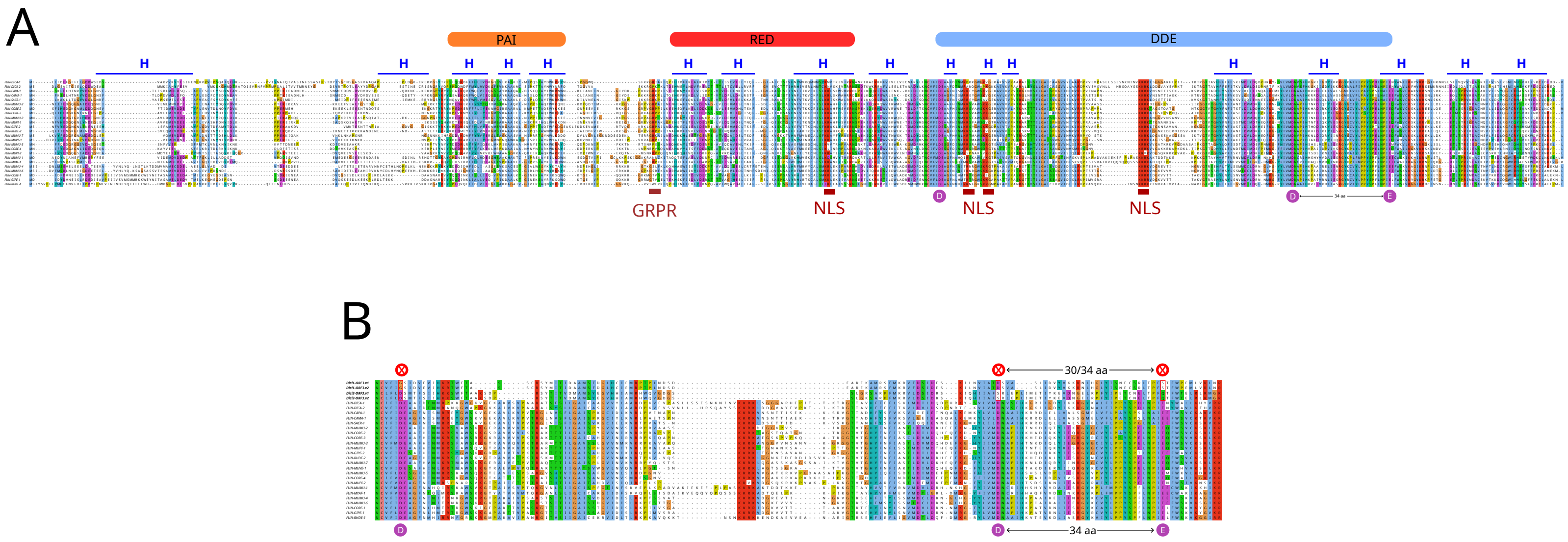

**Supplementary Figure 3.** Characteristics of *Dici1* and *Dici2* transposases. (A) alignment of *Dici* and fungal transposases reveals the main characteristic features, namely a pair of helix-turn-helix (HTH) motifs (PAIRED domain), each consisting of three $\alpha$-helices (blue lines). The GRPR-like motif, located between the two HTH motifs, is particularly well conserved in the top 14 sequences. Four putative nuclear localization signals (NLS), composed of basic amino acids (K or R), were also found to be conserved. The catalytic triad DDE (highlighted with purple circles) is present within the catalytic domain (blue region), with 34 amino acids between the second aspartic acid (D) and glutamic acid (E). The corresponding sequence names, organisms, and accession numbers can be found in Supplementary Table 8; (B) alignment of the four protein sequences of CDS3 (in bold) reveals that it is most likely an inactive remnant of a transposase, as the catalytic triad DDE is not conserved (highlighted in red)

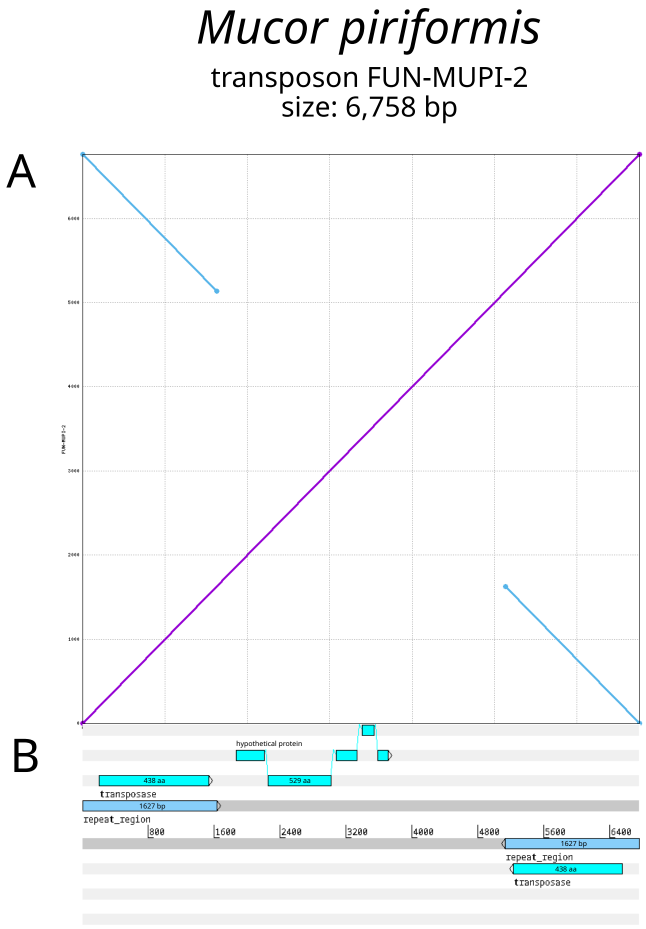

**Supplementary Figure 4.** Characteristics the 6.6-kb transposon detected in *Mucor piriformis*. (A) dotplot obtained from a reciprocal alignment of the complete transposon sequence using MUMmer. The two inverted-repeat regions of 1,627 bp, observed at both extremities, are drawn in blue; (B) the transposon sequence was annotated. The repeat regions contain two identical copies of a CDS that encodes the transposase (438 aa). An additional CDS coding for a hypothetical protein (529 aa) was detected

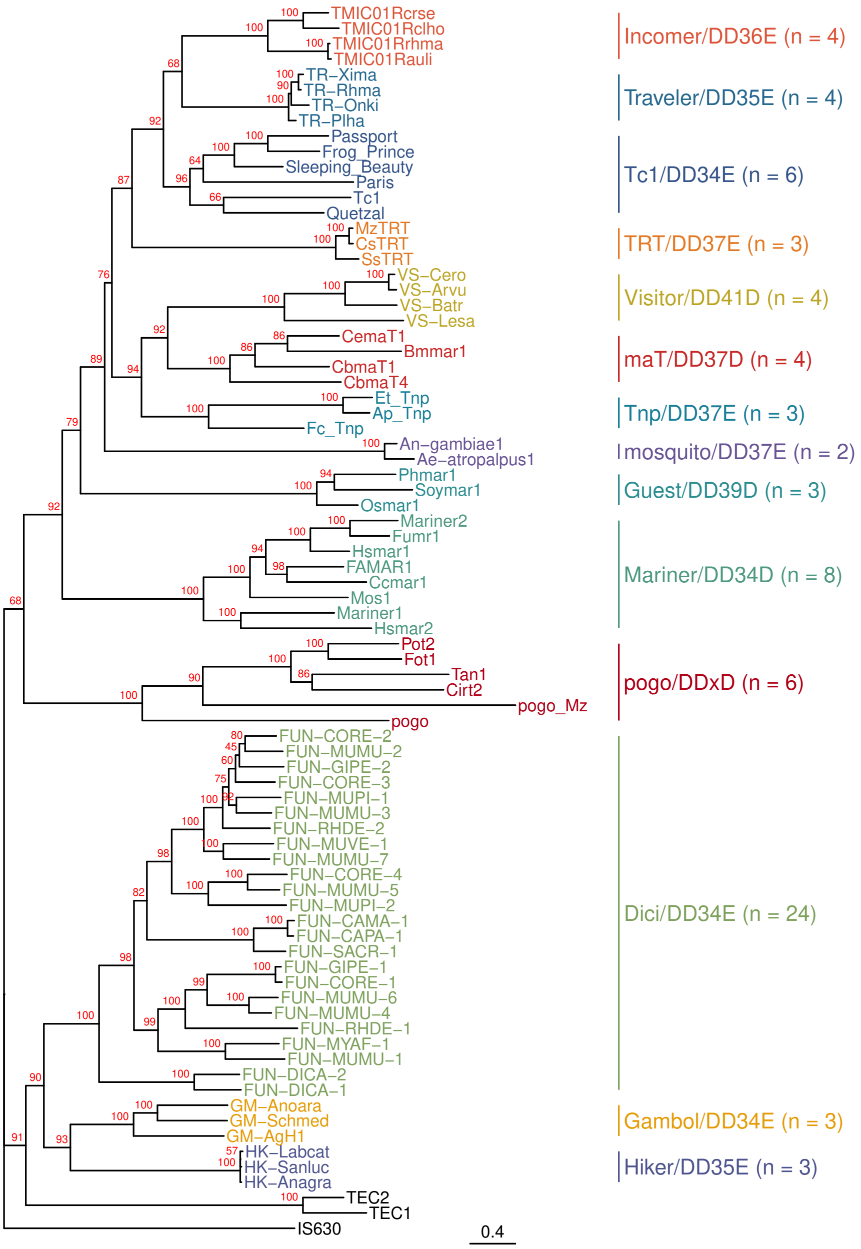

**Supplementary Figure 5.** Phylogenetic tree of the *Dici* transposases alongside transposases belonging to the *IS630-Tc1-mariner* superfamily. The tree was computed using the full-length transposase sequences and a maximum-likelihood method with a bootstrap approach (bootstrap values are shown in red). The *IS630* transposase was used as an outgroup. This tree is identical to the one in the main text, except that the clades are not collapsed. Refer to the Supplementary Table 8 for a detailed description of the transposase name and family

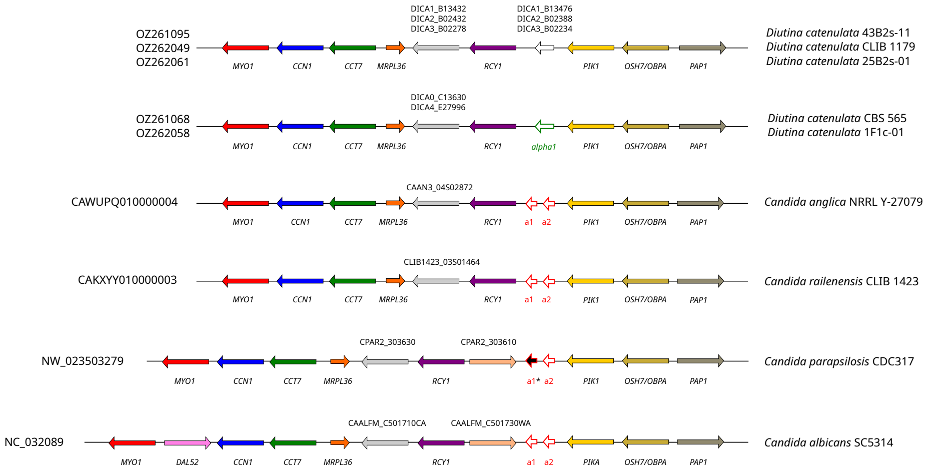

**Supplementary Figure 6.** Structure of the MTL loci in *D. catenulata*, *Candida anglica*, *C. railenensis*, *C. parapsilosis* and *C. albicans*. Species and strain names are on the right, and scaffold or chromosome accession numbers are on the left. Genes in synteny across strains are colored the same and labeled with their corresponding name. Those that encode a protein of unknown function are labeled with their corresponding locus tag. Pseudogenes are marked with an asterisk

#### 4 Supplementary tables

Additional spreadsheet documents (XLSX)

- **Supplementary Table 1.** *D. catenulata* strains used in the study. Also includes accession numbers for sequencing and assembled genome data.
- **Supplementary Table 2.** Genetic diversity and differentiation statistics measured in *D. catenulata*.
- **Supplementary Table 3.** Assembly and annotation statistics of *D. catenulata* assembled genomes.
- **Supplementary Table 4.** Genes associated with drug resistance and virulence detected in *D. catenulata*.
- **Supplementary Table 5.** Characteristics and chromosomal positions of the RNA transposons discovered in *D. catenulata* genomes.
- **Supplementary Table 6.** Characteristics and chromosomal positions of the *Dici* transposons discovered in *D. catenulata* genomes.
- **Supplementary Table 7.** List of synteny blocks detected between the *D. catenulata* CBS 565 genome and those of the other four cheese strains.
- **Supplementary Table 8.** Characteristics of the *Dici* transposons newly identified in Fungi. Also contains the list of the *IS630-Tc1-mariner* transposase sequences used to generate the phylogenetic tree.
- **Supplementary Table 9.** Sequencing statistics for the five *D. catenulata* assembled genomes. Also contains short-reads mapping statistics obtained for the 61 *D. catenulata* stains.
